# Placental and Immune Cell DNA Methylation Reference Panel for Bulk Tissue Cell Composition Estimation in Epidemiological Studies

**DOI:** 10.1101/2024.05.06.588886

**Authors:** Kyle A. Campbell, Justin A. Colacino, John Dou, Dana C. Dolinoy, Sung Kyun Park, Rita Loch-Caruso, Vasantha Padmanabhan, Kelly M. Bakulski

## Abstract

To distinguish DNA methylation (DNAm) from cell proportion changes in whole placental tissue research, we developed a robust cell type-specific DNAm reference to estimate cell composition. We collated newly collected and existing cell type DNAm profiles quantified via Illumina EPIC or 450k microarrays. To estimate cell composition, we deconvoluted whole placental samples (n=36) with robust partial correlation based on the top 50 hyper- and hypomethylated sites per cell type. To test deconvolution performance, we evaluated RMSE in predicting principal component one of DNAm variation in 204 external placental samples. We analyzed DNAm profiles (n=368,435 sites) from 12 cell types: cytotrophoblasts (n=18), endothelial cells (n=19), Hofbauer cells (n=26), stromal cells (n=21), syncytiotrophoblasts (n=4), six lymphocyte types (n=36), and nucleated red blood cells (n=11). Median cell composition was consistent with placental biology: 60.4% syncytiotrophoblast, 17.1% stromal, 8.8% endothelial, 4.5% cytotrophoblast, 3.9% Hofbauer, 1.7% nucleated red blood cells, and 1.2% neutrophils. Our expanded reference outperformed an existing reference in predicting DNAm variation (15.4% variance explained, IQR=21.61) with cell composition estimates (RMSE:10.51 vs. 11.43, p-value<0.001). This cell type reference can robustly estimate cell composition from whole placental DNAm data to detect important cell types, reveal biological mechanisms, and improve casual inference.

## Introduction

Prenatal environmental or other exposures during gestation may induce perturbations to placental epigenetic programming, including DNA methylation, that ultimately result in disease [1]. Epigenetic dysregulation is characteristic of diseases such as cancer [2], neurodegeneration [3], and cardiovascular disease [4]. A recent review identified that between 2016 and 2021, 28 studies linked air pollution, chemical, metals, psychosocial, and smoking exposures to the placental epigenome [5]. A critical limitation of these studies, however, is a failure to account for cell type heterogeneity [5, 6]. In fact, two studies identified an association between prenatal mercury exposure and DNA methylation in fetal cord blood as well as a shift in cord blood cell type proportions, precluding biological interpretation as to the causal factor [7, 8]. Bioinformatic reference-based deconvolution has emerged as a promising strategy to account for cell heterogeneity in bulk tissue assays by estimating cell type proportions from previously collected cell type-specific molecular profiles [9–11]. However, placental cell type-specific DNA methylation references are limited [12]. Several studies have individually profiled DNA methylation of cytotrophoblasts and Hofbauer cells at term [13, 14]. One study conducted whole genome DNA methylation characterization of cytotrophoblasts from two term placentas [14]. DNA methylation from term Hofbauer cells was characterized on the Illumina Infinium Human Methylation27 BeadChip microarray [13]. This platform measured DNA methylation at ∼27,000 sites across the genome, which is much lower coverage than the ∼850,000 sites now available on the EPIC version of this platform. Only recently has one study profiled trophoblasts, stromal, endothelial cells, and Hofbauer cells from fractions of placental cells enriched using fluorescence-activated cell sorting, and a separate filter-enriched syncytiotrophoblast fraction on the Illumina EPIC DNA methylation microarray [15]. One caveat is the potential for maternal peripheral immune cells and the highly epigenetically distinct nucleated red blood cells present only in fetal cord blood to be intermingled with placental cell types in villous tissue samples [16]. Indeed, a DNA methylation epigenome-wide association study in fetal cord blood found that nucleated red blood cells explained most of the association between gestational age and DNA methylation [17]. A robust placental deconvolution reference requires additional placental cell type-specific DNA methylation profiles and must account for other epigenetically distinct cell types present at the fetal-maternal interface.

This study aims to address the current limitations of existing approaches to account for cellular heterogeneity in DNA methylation studies of the human placenta. Here, we generate primary cell type-specific placental DNA methylation profiles for major placental cell types isolated via magnetic bead activated cell sorting and integrate the results with existing placental, adult peripheral immune cell, and nucleated red blood DNA methylation profiles to create a unified placental DNA methylation deconvolution panel for reference-based deconvolution. Placental cell type-specific genome-wide DNA methylation panels can then be implemented in numerous existing or future studies that utilize placental villous tissue samples at term to improve precision, account for non-placental cell types such as maternal peripheral immune cells, reduce sources of potential bias due to cell type heterogeneity, and improve biological inference.

## Methods

### Placental tissue sample collection and dissociation

We collected placentas shortly after delivery from healthy, full term, singleton uncomplicated Cesarean section deliveries at the University of Michigan Von Voigtlander Women’s Hospital between February 2019 and March 2020. All study participants provided informed consent. This study protocol was approved by the University of Michigan Institutional Review Board (#HUM00017941). Villous placental tissue samples were collected, washed in phosphate buffered saline (PBS, Invitrogen #10010049), blotted dry, and minced from the maternal-facing side of placenta after trimming away the basal and chorionic plates and scraping tissue from blood vessels until at least 10 g of villous tissue were collected. Approximately 200 mg of villous tissue was set aside and snap-frozen in liquid nitrogen.

We subjected isolated fetal villous tissue to three sequential enzymatic digestions followed by discontinuous density gradients using slight modifications on established methods to isolate cytotrophoblasts, fibroblasts, and Hofbauer cells [18, 19]. Enzymatic digestions were carried out in a Trypsin-DNase I solution containing 0.25% trypsin (Corning, #25-050-CI), 0.20% DNase I (Roche, #10104159001), 25 mM N-2-hydroxyethylpiperazine-N’-2-ethanesulfonic acid (HEPES, Millipore Sigma, H4034-25G), 2.0 mM CaCl_2_ (Millipore Sigma, #C7902-500G), and 0.8 mM MgSO_4_ (Millipore Sigma, M2643-500G) in Hanks’ balanced salt solution (HBSS, Invitrogen #14185052) at 37°C with shaking at 150 rpm in a large shaker bath [18]. The first digestion lasted 15 minutes and the partially digested villous tissue was strained through a 104 μm mesh with a pestle (Millipore Sigma, #CD1-1KT) and rinsed with 3 mL heat-inactivated fetal bovine serum (FBS, Corning, # MT35016CV) and 2 rounds of 10 mL DF medium, consisting of 10% FBS, 1% antibiotic-antimycotic (Invitrogen, #15240062), in Dulbecco’s Modified Eagle Medium/Ham’s F-12 (DMEM/F12 medium, Invitrogen, #12634010) to quench 15 mL of Trypsin-DNase I digestion solution and facilitate straining.

The second and third digests were performed for 30 minutes each but were otherwise identical to the first digestion. The supernatant from each digest was filtered sequentially through another 100 μm filter and a 70 μm filter. PBS washes were used as needed to facilitate straining. Adapting a previously published protocol [20], the 70 μm filters were immediately inverted into a sterile petri dish and washed with DF medium to produce a syncytiotrophoblast-enriched fraction that was pooled from each of the three digests and cryopreserved in a cryopreservation solution containing 10% dimethyl sulfoxide (DMSO, Millipore Sigma #D8418) in FBS in liquid nitrogen. The supernatant from the second and third digests were combined and loaded onto a 30%/35%/45%/50% Percoll discontinuous gradient consisting of 90% Percoll (GE Healthcare, #17089101) mixed with 25 mM HEPES in HBSS in the appropriate concentrations. We performed density gradient centrifugation by centrifuging the column at 1000 rcf for 20 minutes at room temperature with no brake [18]. To create a cytotrophoblast-enriched fraction, we collected the cell layer between the 35% and 45% fraction, which was cryopreserved in 10% DMSO in FBS in liquid nitrogen [18].

To isolate a Hofbauer cell and fibroblast-enriched fraction, we subjected the tissue to a final 1 mg/mL Collagenase A, 0.2 mg/mL DNase I 20 mL enzymatic digestion in an RPMI medium containing 25 mM HEPES, 10% FBS, and 1% antibiotic/antimycotic for 1 hour at 37 °C. The digestion supernatant was strained through a 104 μm mesh with a pestle (Millipore Sigma, #CD1-1KT) and rinsed with 3 mL FBS and 2 rounds of 10 mL RPMI medium to quench 20 mL of Collagenase A-DNase I digestion solution and facilitate straining. We performed sequential density gradient centrifugations by centrifuging columns at 1200 rcf for 20 minutes at room temperature with no brake. We loaded the single cell suspension onto a 20%/40% Percoll discontinuous gradient and collected cells from the 20%/40% interface. We loaded these cells onto a second 20%/25%/30%/35% discontinuous Percoll gradient. Cells were collected and pooled from the 20%/25%, 25%/30%, and 30%/35% interfaces. These cells were cryopreserved in 10% DMSO in FBS in liquid nitrogen.

### Magnetic activated bead cell type sorting

A schematic overview of the magnetic activated bead sorting protocol is presented as **Supplementary Figure 1**. Cryopreserved single cell suspensions were quickly thawed at 37 °C, size-filtered at 30 µm to eliminate debris, and resuspended in 4 °C DF medium for the cytotrophoblast fraction or RPMI medium for the Hofbauer cell/fibroblast fraction. Both fractions were subsequently resuspended in a 4 °C BSA rinsing buffer, consisting of 0.5% Bovine Serum Albumin stock solution (Miltenyi Biotec, #130-091-376) in autoMACS washing solution (Miltenyi Biotec, #130-092-987). All media and cell suspensions were kept on ice for the remainder of the magnetic-activated sorting. Following manufacturer’s instructions, the Miltenyi Biotec octoMACS platform (Miltenyi Biotec, #130-042-108) was used to magnetically sort the target placental cell types cytotrophoblasts, Hofbauer cells, and fibroblasts. Briefly, the cytotrophoblast fraction was incubated for 10 minutes at 4 °C in the dark with 1:11 REAfinity™ allophycocyanin (APC)-conjugated anti-human leukocyte antigen (HLA)-ABC antibody (APC-anti-HLA-ABC, Miltenyi Biotec, #130-101-467, lot #5191002371). The Hofbauer/fibroblast fraction was incubated for 10 minutes at 4 °C in the dark with 1:11 REAfinity™ allophycocyanin-conjugated anti-epidermal growth factor (EGFR) antibody (APC-anti-EGFR, Miltenyi Biotec, #130-110-529, lot #5190107002). Each fraction was washed with BSA rinsing buffer and incubated for 15 minutes at 4 °C in the dark with 1:5 anti-allophycocyanin MACS® MicroBeads (Miltenyi Biotec, #130-090-855, lot #5190809239). Each fraction was loaded onto an MS column (Miltenyi Biotec, #130-042-201) for magnetic separation. The EGFR-Hofbauer/fibroblast fraction was washed and incubated for 10 minutes at 4 °C in the dark with 1:11 REAfinity™ phycoerythrin (PE)-conjugated anti-CD10 antibody (anti-CD10-PE, Miltenyi Biotec, #130-114-5-2, lot #5190109168). The Hofbauer/fibroblast fraction was then washed and incubated for 15 minutes at 4 °C in the dark with 1:5 anti-phycoerythrin MACS® MicroBeads (Miltenyi Biotec, #130-105-639). The Hofbauer/fibroblast fraction was loaded onto an MS column for magnetic separation. The CD10-fraction is enriched for fibroblasts and the CD10+ fraction is enriched for Hofbauer cells [18]. In downstream, integrative analyses, our fibroblast fraction is labelled stromal to comport with the cell type labels used in secondary analysis studies.

### DNA Extraction

To collect whole tissue DNA samples, approximately 15mg of snap-frozen villous tissue was added to 180 μL Buffer ATL (Qiagen, #939011) and 20 μL proteinase K (Qiagen, #19131) to Lysing Matrix D vials (MP Biomedicals, #116913050-CF). Samples were homogenized on the MP-24 FastPrep homogenizer (MP Biomedicals, #116004500) at 6m/s, setting MP24x2 for 35 seconds. The cryogenically stored cell type fractions were quickly thawed at 37 °C for less than five minutes. DNA was extracted from each fraction using the DNeasy Blood and Tissue Kit (Qiagen, #69504, lot #157020715, 1630203096, 563011175) according to manufacturer’s instruction for each magnetically separated sample, the syncytiotrophoblast-enriched sample, and the bulk whole villous tissue sample after homogenization.

### DNA methylation measurement

DNA was submitted to the University of Michigan Epigenomics Core for quality assessment and bisulfite conversion. DNA quantity was measured with the Qubit High Sensitivity dsDNA assay (ThermoFisher Scientific, #Q32854). DNA quality was assessed with the TapeStation genomic DNA kit (Agilent, #5067-5365). Samples too diluted to be compatible with the kit were first concentrated using AMPure Beads (Beckman Coulter, #A63880) and re-quantified. An aliquot of 250 ng DNA from each sample was bisulfite converted using the EZ DNA Methylation kit (Zymo Research, #D5001) according to manufacturer’s instructions for Illumina DNA methylation arrays. Samples were then sent to the University of Michigan Advanced Genomics Core for hybridization to the Infinium MethylationEPIC BeadChip v1.0 array (Illumina, #WG-317-1003), washing, and scanning according to the manufacturer’s instructions. Samples were randomized across columns and rows (within plates) to reduce potential confounding due to batch effects.

### DNA methylation data preprocessing and quality control

Data were processed in R statistical software (version 4.2.2). In addition to the data generated in this study, we also downloaded data from three previously published cell type-specific DNA methylation studies: 1) adult blood immune cells (Gene Expression Ominbus (GEO) accession GSE110554, accessed via FlowSorted.Blood.EPIC package v1.99.5 [21]), including B cells, CD4+ T cells, CD8+ T cells, monocytes, natural killer cells, and neutrophils [22]; 2) umbilical cord blood (accessed via FlowSorted.CordBloodCombined.450k package v1.8.0 [23]) [16], including a combined dataset of nucleated red blood cell samples from Bakulski et al. [24] and de Goede et al. [25]; and 3) placental cell types (GEO accession GSE159526, accessed via ExperimentHub package v2.0.0 [26]) [15] endothelial, Hofbauer, stromal, trophoblast, and syncytiotrophoblast cell types as well as whole tissue placental samples. Raw DNA methylation data were preprocessed and managed via the minfi package v1.38.0 [27] and the ewastools package v1.7 [28].

Datasets generated on the Illumina 450k microarray were preprocessed and quality controlled separately from those generated on the Illumina EPIC DNA methylation microarray. We preprocessed raw DNA methylation data with Noob background correction and dye-bias normalization. Sample and probe inclusion and exclusion was visualized using a flow chart. No samples were flagged for low-intensity values below 10.5 relative fluorescence units or for genotype or sex mismatches. For the EPIC array samples (n=189), we excluded 1 probe with an average bead count less than 5, 19,760 probes with >5% failure rate for detection p-value >0.01, 23,448 cross-reactive probes [29], and 17,767 sex-specific probes. For the 450k array samples (n=132), we excluded 20,024 probes with >5% failure rate for detection p-value >0.01, 14,151 cross-reactive probes [29], and 9,646 sex-specific probes. Bead hybridization information was unavailable for the 450k umbilical cord samples and only the nucleated red blood cells (n=11) were retained for downstream analysis. Finally, to correct for type I vs. type II probe bias, we also performed Beta MIxture Quantile dilation normalization [30] via the ChAMP package v2.22.0 [31]. All downstream DNA methylation analyses used the methylation rate for a given site, also known as beta values.

To make our results generalizable to the Illumina 450k array, we subset the EPIC array samples to only those DNA methylation sites shared across both arrays for all downstream analyses. To assess cell fraction purity, we assessed unsupervised DNA methylation profile clustering and predicted cell type lineage from an external reference [32]. To reduce dimensionality for unsupervised DNA methylation clustering, we selected the top 10% highest variance DNA methylation sites and performed a principal component analysis. Iterative clustering and sub-clustering with the Seurat package v4.0.1 [33] function FindClusters at different resolution parameters were evaluated using cluster stability via clustering trees with clustree package v0.4.4 [34]. We compared data from cell fractions across study sources by visualizing clustering results with uniform manifold projection [35]. Samples that clustered inconsistently across placental datasets were removed from downstream analyses as suspected lower cell type purity samples. As an independent measure of sample purity, we predicted the lineage composition (epithelial, immune cell, or stromal) of each sample with the robust partial correlation algorithm and the centEpiFibIC.m dataset [32] implemented in EpiDISH package v2.8.0 [36]. We compared the distribution of lineage composition between included and excluded cell type samples with the nonparametric Mann-Whitney test with an alternative hypothesis that the appropriate cell type lineage would be higher in the included sample compared to the excluded. For all included samples, we used a table to describe the number and frequency of samples by study source, fetal sex, cell type, Illumina array type, and cell separation method.

### Characterization of cell type fractions with differential methylation analysis and biological process enrichment

We excluded biologic replicate whole tissue samples for cell type characterization and deconvolution reference analyses. To characterize cell type fractions, we assessed cell type-specific DNA methylation patterns by fitting linear models adjusted for sex with empirical Bayes standard error moderation in the limma package v3.48.3 [37]. To identify cell type-specific differentially methylated sites, we compared percent methylation values in one cell type against the average across other cell types. We used false discovery adjusted q-values to account for multiple comparisons. We visualized differences in DNA methylation between cell types using volcano plots of the average methylation difference and the -log_10_(q-value). To be considered biologically meaningful, we instituted an absolute percent methylation difference threshold cutoff of 10%. Statistical significance was assessed at a false discovery adjusted q-value < 0.001. We visualized overlap in significant sites across cell types with an upset plot, separately for hyper- and hypomethylated sites. Annotations for biological interpretation of differential methylation results were retrieved from GEO Platform Accession GPL13534 [38], NCBI Reference Sequence [39], The Genome Browser at University of California Santa Cruz [40], ENCODE [41], and GeneHancer [42].

To characterize the biological significance of differential methylation results, we performed gene ontology enrichment for the top 10,000 differentially methylated sites ranked by descending absolute value of the model contrast test statistic among sites that met biological effect size and statistical significance thresholds per cell type with the missMethyl package v1.26.1 [43]. This approach identifies overrepresented genes and their ontologies among differentially methylated sites while accounting for the number of sites tested per gene as well as probes associated with multiple genes. We visualized overlap in enriched gene ontologies using an upset plot.

### Creation and application of a placental DNA methylation deconvolution reference

To identify cell type-discriminating DNA methylation sites, we ranked the top 50 each of hyper- and hypomethylated sites per cell type by F-test using the minfi package v1.38.0 [27]. At these probes, we visualized DNA methylation levels across cell types using a heat map. These cell type-discriminating DNA methylation sites were used as the cell type DNA methylation references for deconvolution of whole tissue placental villous samples present in the analyzed datasets. We used the robust partial correlation algorithm implemented in EpiDISH package v2.8.0 [36] to perform deconvolution. We visualized estimated cell proportions in whole tissue using a scatter plot and calculated the estimated proportion quartiles.

### Comparative assessment of placental DNA methylation deconvolution reference performance

To assess the performance of our placental DNA methylation deconvolution reference, we compared its ability to predict DNA methylation variation [44] in a preprocessed placental whole tissue Illumina EPIC microarray DNA methylation dataset comprised of 204 placentas across three cohorts (GEO accession number GSE232778) [45] against the only existing multi-cell type reference [15]. To capture a low-dimensional representation of placental DNA methylation variation, we performed a principal component analysis using prcomp v4.2.2 [46] of site-specific methylation rates in GSE232778 and extracted the first principal component. To calculate GSE232778 cell composition estimates, we deconvoluted placental cell composition using the robust partial correlation algorithm implemented in EpiDISH package v2.8.0 [36] with either our updated reference or the existing reference [15]. To directly compare our updated reference to the existing reference in the planet v1.6.0 package [15], we evaluated which model yielded a smaller root mean square error of prediction (RMSEP) across 1,000 trials of 10-fold cross-validation for linear regression models using xvalglms v0.1.8 [47]. These models used the first principal component of GSE232788 placental DNA methylation variation as the response variable and either our updated references’ cell composition deconvolution estimates, or the existing references’ estimates as the predictor variables.

## Results

### Cell type sorting captures placental and immune cell type-specific DNA methylation profiles

We analyzed the placental datasets together with DNA methylation profiles from adult blood peripheral immune cells and umbilical cord nucleated red blood cells, which may also be present in placental samples, especially those collected in large epidemiologic or biobanking studies. We performed several quantitative and qualitative tests to assess cell type fraction purity. Consistent with successful cell type isolation, unsupervised clustering revealed that DNA methylation samples clustered by their target cell type and UMAP projection shows cell type lineages colocalize (**Figure 1**). We excluded 37 placental cell type samples that clustered inconsistently with their primary cell type cluster: 25 cell type samples from this study (5 cytotrophoblast, 1 Hofbauer, 11 stromal, and 8 syncytiotrophoblast samples; and 12 placental cell type samples from Yuan et al. [15]: 11 cytotrophoblast, and 1 syncytiotrophoblast sample) (**Supplementary Figure 2** Uniform manifold approximation and projection (UMAP) of DNA methylation data showing how cell type samples that clustered inconsistently with their primary cluster were excluded from downstream analysis.. Samples who To confirm cell type sample purity, we estimated broad cell type lineage (epithelial, stromal, or immune cell) from an independent deconvolution reference [32] and compared the appropriate estimated lineage composition between included and excluded samples. Included versus excluded epithelial composition for cytotrophoblasts was 84.5% vs. 55.6% (p<0.001) and 80.5% vs. 43.3% for syncytiotrophoblast (p<0.001); fibroblast composition for stromal cells was 74.0% vs. 16.3% (p<0.001); and immune cell composition was 78.4% vs. 43.0% (p=0.037) for Hofbauer cells.

**Figure 1.**
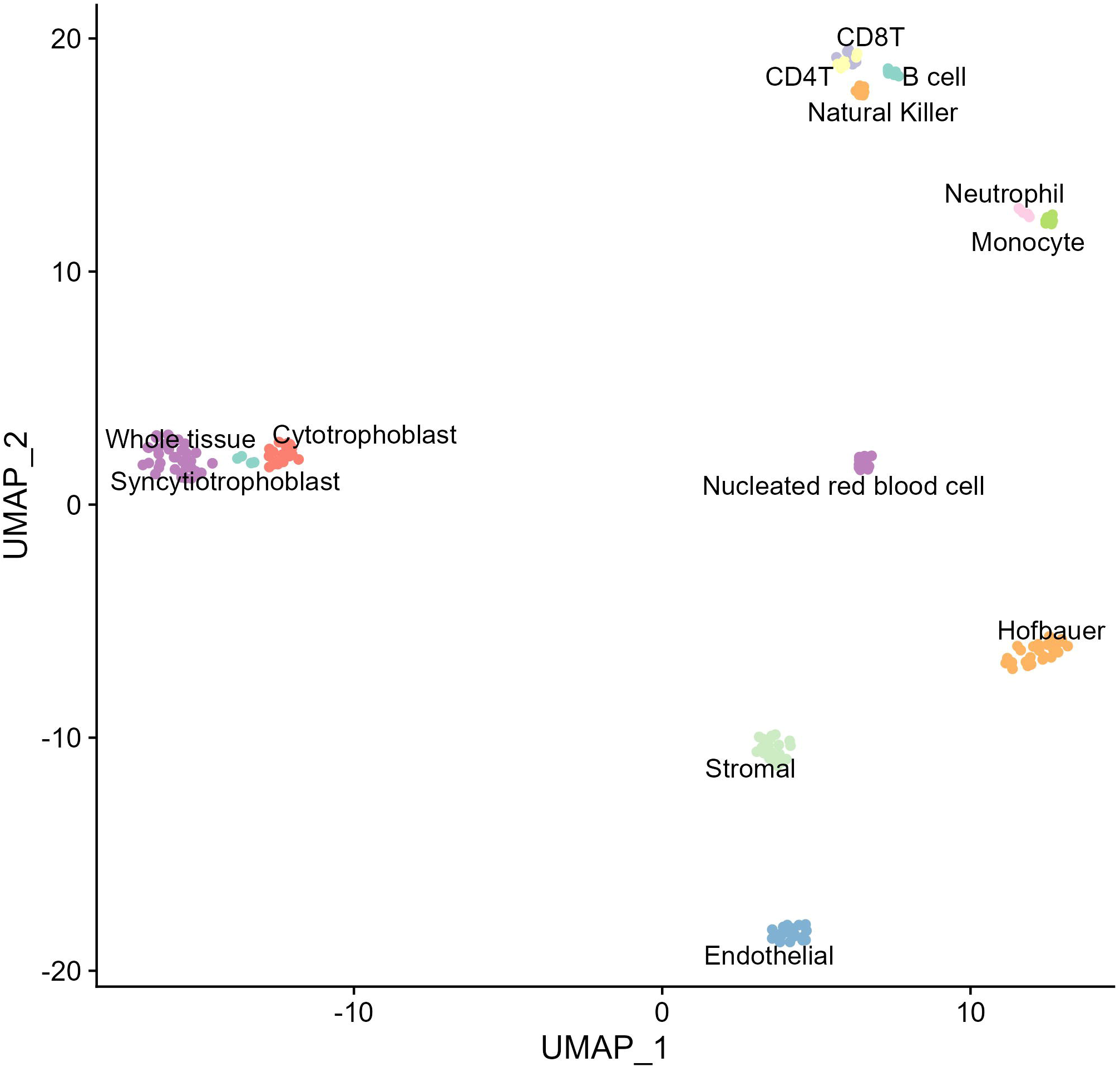
Low-dimensional representation of variable DNA methylation sites across all placental and immune cell types and whole placental tissue samples shows samples cluster by cell type.

After quality control procedures (**Supplementary Figure 3**), we analyzed DNA methylation data across 368,435 sites from 135 new and previously published DNA methylation profiles from 12 cell type-specific fractions, including cytotrophoblasts (n=18), endothelial cells (n=19), Hofbauer cells (n=26), stromal cells (n=21), syncytiotrophoblast-enriched fractions (n=4) [15], six types of adult lymphocytes (n=36) [22], and nucleated red blood cells (n=11) [16], as well as 36 bulk placental tissue samples (**Table 1**).

**Table 1.** Analytic table of DNA methylation samples used for deconvolution reference.

| Characteristic | Campbell, N = 37<sup>†</sup> | Yuan, N = 87<sup>†</sup> | Salas, N = 36<sup>†</sup> | Bakulski, N = 4<sup>†</sup> | de Goede, N = 7<sup>†</sup> |
| --- | --- | --- | --- | --- | --- |
| **Fetal Sex** |  |  |  |  |  |
| Female | 24 (65%) | 48 (55%) | 6 (17%) | 3 (75%) | 5 (71%) |
| Male | 13 (35%) | 39 (45%) | 30 (83%) | 1 (25%) | 2 (29%) |
| **Cell Type** |  |  |  |  |  |
| B cell | 0 (0%) | 0 (0%) | 6 (17%) | 0 (0%) | 0 (0%) |
| CD4T | 0 (0%) | 0 (0%) | 6 (17%) | 0 (0%) | 0 (0%) |
| CD8T | 0 (0%) | 0 (0%) | 6 (17%) | 0 (0%) | 0 (0%) |
| Cytotrophoblast | 10 (27%) | 8 (9.2%) | 0 (0%) | 0 (0%) | 0 (0%) |
| Endothelial | 0 (0%) | 19 (22%) | 0 (0%) | 0 (0%) | 0 (0%) |
| Hofbauer | 8 (22%) | 18 (21%) | 0 (0%) | 0 (0%) | 0 (0%) |
| Monocyte | 0 (0%) | 0 (0%) | 6 (17%) | 0 (0%) | 0 (0%) |
| Neutrophil | 0 (0%) | 0 (0%) | 6 (17%) | 0 (0%) | 0 (0%) |
| Natural Killer | 0 (0%) | 0 (0%) | 6 (17%) | 0 (0%) | 0 (0%) |
| Nucleated red blood cell | 0 (0%) | 0 (0%) | 0 (0%) | 4 (100%) | 7 (100%) |
| Stromal | 2 (5.4%) | 19 (22%) | 0 (0%) | 0 (0%) | 0 (0%) |
| Syncytiotrophoblast | 0 (0%) | 4 (4.6%) | 0 (0%) | 0 (0%) | 0 (0%) |
| Whole tissue | 17 (46%) | 19 (22%) | 0 (0%) | 0 (0%) | 0 (0%) |
| **Illumina Array Type** |  |  |  |  |  |
| 450k | 0 (0%) | 0 (0%) | 0 (0%) | 4 (100%) | 7 (100%) |
| EPIC | 37 (100%) | 87 (100%) | 36 (100%) | 0 (0%) | 0 (0%) |
| **Cell Separation Method** |  |  |  |  |  |
| Fluorescence activated cell sort | 0 (0%) | 87 (100%) | 36 (100%) | 0 (0%) | 7 (100%) |
| Magnetic activated cell sort | 37 (100%) | 0 (0%) | 0 (0%) | 4 (100%) | 0 (0%) |

| Characteristic | Campbell, N = 37<sup>1</sup> | Yuan, N = 87<sup>1</sup> | Salas, N = 36<sup>1</sup> | Bakulski, N = 4<sup>1</sup> | de Goede, N = 7<sup>1</sup> |
| --- | --- | --- | --- | --- | --- |
| <sup>1</sup>n (%) |  |  |  |  |  |

### Placental and immune cell type-specific DNA methylation patterns

To identify placental and immune cell type-defining DNA methylation patterns and provide additional evidence of successful cell type isolation, we compared site-specific percent DNA methylation in one cell type against the average methylation rate across all other cell types (**Supplementary Table 1**). Consistent with successful cell type separation, we observed large-scale, genome-wide differential DNA methylation patterns across cell types. Of placental cell types, the number of hypermethylated sites ranged from 41,005 in endothelial cells to 59,972 in cytotrophoblasts. Among placental cell types, when compared to average methylation across all other cell types, the number of hypomethylated sites per placental cell type ranged from 44,544 in Hofbauer cells to 89,846 in cytotrophoblasts (**Figure 2**). Nucleated red blood cells had the largest number of differentially methylated sites at 259,414. Of placental cell types, cytotrophoblasts had the largest number of differentially methylated sites at 149,818, followed by syncytiotrophoblasts at 141,686.

**Figure 2.**
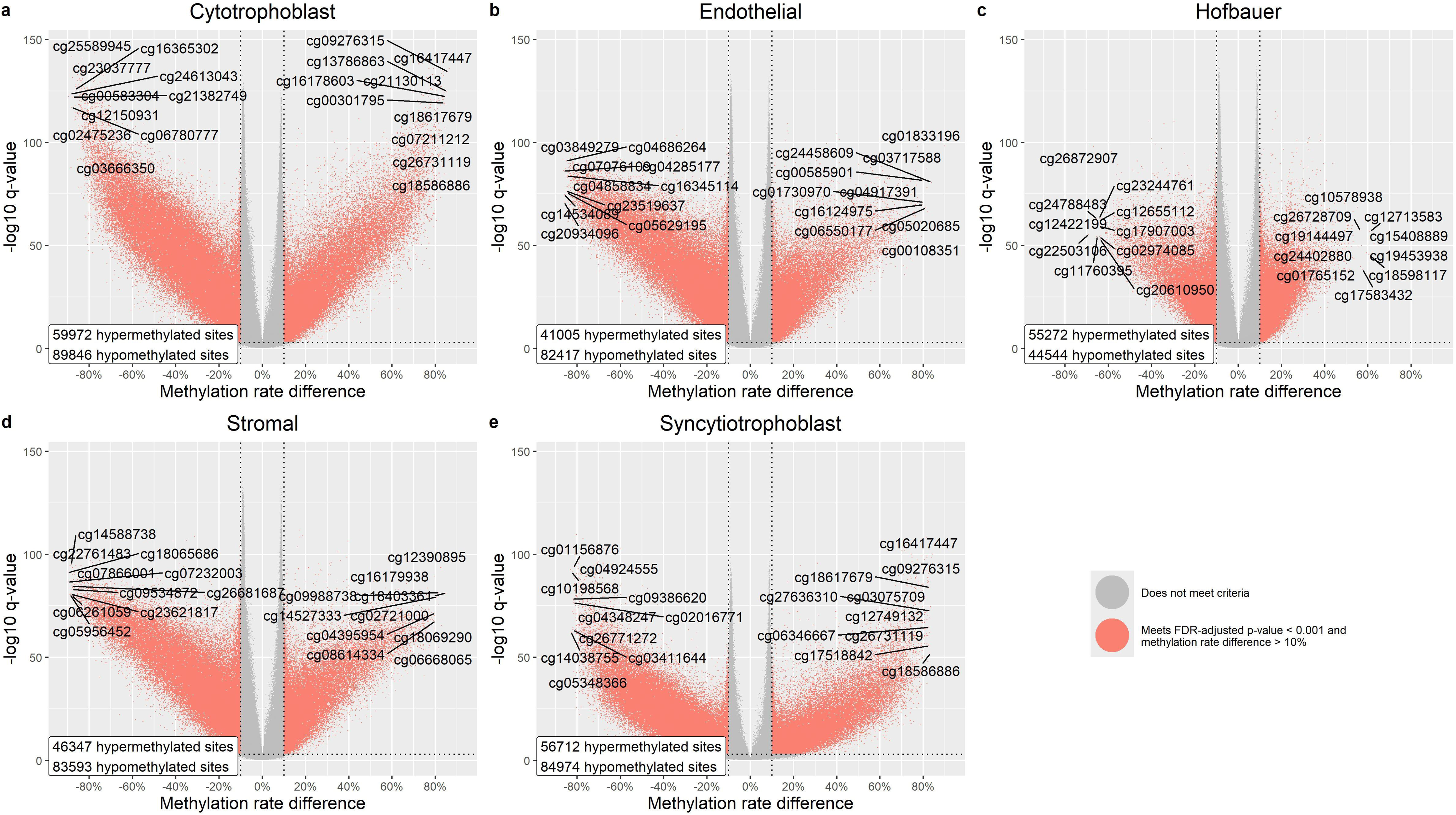
Placental cell type differential DNA methylation. Volcano plots comparing site-specific DNA methylation levels in one cell type against the average across all other cell types.

Examination of top differentially methylated hits specific to each of the five placental cell types (cytotrophoblast, endothelial, Hofbauer, stromal, and syncytiotrophoblast) and nucleated red blood cells revealed highly specific DNA methylation patterns with biologically relevant gene annotations (**Figure 3**). For example, cytotrophoblast samples exhibited hypomethylation at cg23757825 compared to other cell types (−85.2% average methylation difference, p_adj_<0.001), the target of an enhancer and a site annotated to the gene body of epidermal growth factor receptor (*EGFR*), a protein used to isolate cytotrophoblasts [15, 18]. Endothelial samples were strongly hypomethylated at cg00097800 (−83.3%, p_adj_<0.001), annotated to exon one of the oxygen-sensing transcription factor *EPAS1/HIF2a*, a gene critical to the migration, growth, and differentiation of endothelial cells [48]. Hofbauer samples were hypomethylated at cg06713769 (−48.5%, p_adj_<0.001), annotated to *LYVE-1*, a protein previously identified in Hofbauer cells [49]. Stromal samples exhibited hypomethylation at cg26574240 (−83.7%, p_adj_<0.001), annotated as within 200 base pairs of a transcription start site for *LMOD3*, a protein responsible for actin polymerization [50]. Syncytiotrophoblast samples were hypomethylated at cg19368383 (−79.6%, p_adj_<0.001), annotated to the body of *HPS1*, a protein essential to the construction of lysosome-related organelles [39]. Nucleated red blood cells exhibited extreme hypomethylation at cg08985539 (−97.8%, p_adj_<0.001), annotated to the body of *POLN*, encoding DNA polymerase Nu [39].

**Figure 3.**
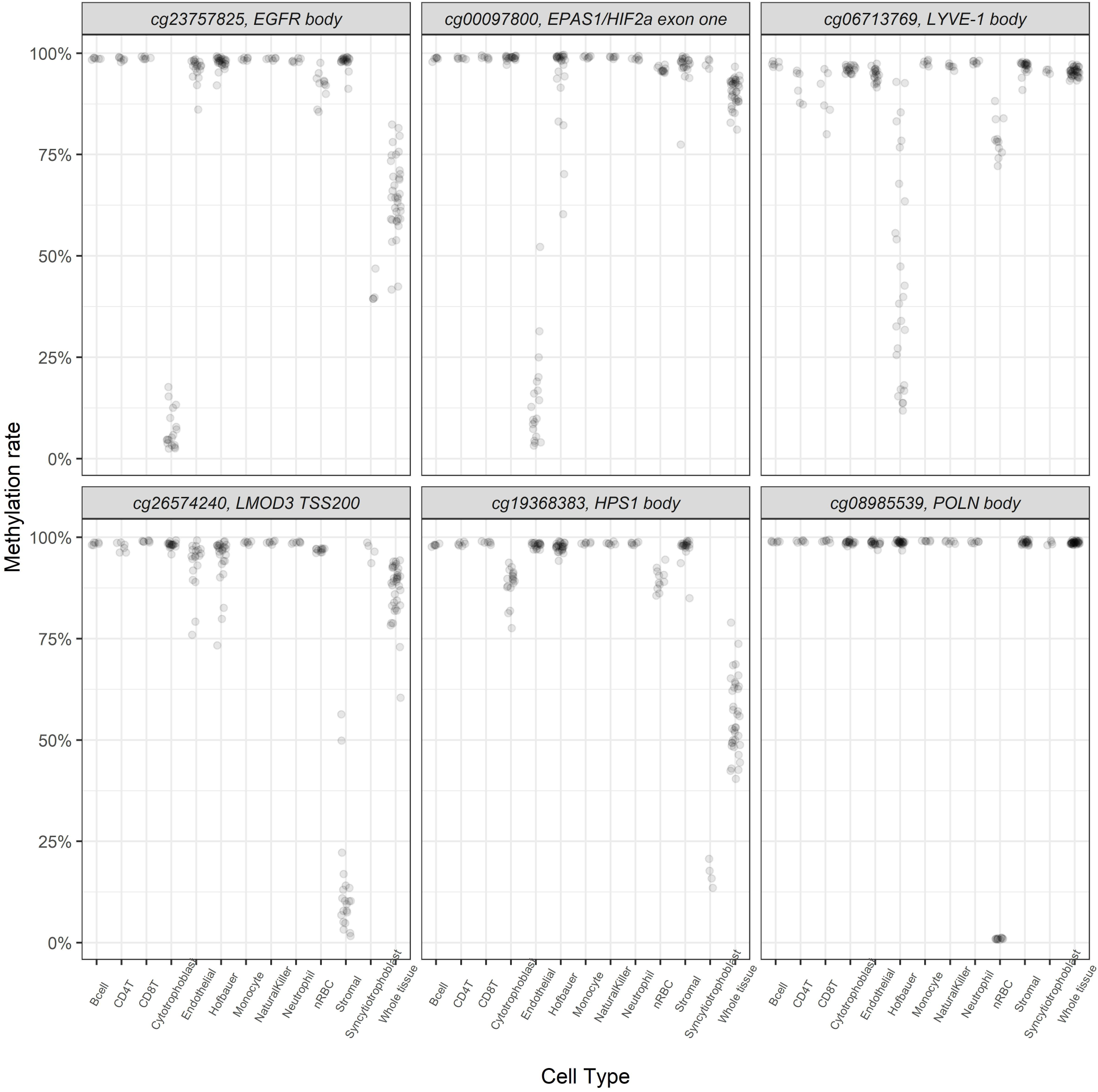
Top differentially methylated sites unique to the five placental cell types (cytotrophoblast, endothelial, Hofbauer, stromal, and syncytiotrophoblast) and nucleated red blood cells with gene annotations reveal highly specific DNA methylation profiles by cell type.

### Placental biologic processes revealed at differentially methylated sites

To identify the functional biological relevance of differential methylation results, we performed biologic process ontology enrichment of differentially methylated genes (**Supplementary Table 2**). The number of biological processes unique to placental cell types ranged from 217 for endothelial cells to 7 for cytotrophoblast. Biological process ontology enrichment highlighted cell type-specific functions consistent with successful cell type isolation. For example, in syncytiotrophoblasts, actin filament-based process (p_adj_<0.001), extracellular matrix organization (p_adj_=0.002), and sialyation (p_adj_=0.008) were enriched among genes differentially methylated. Differentially methylated cytotrophoblast genes were enriched for external encapsulating structure organization (p_adj_=0.026), cell adhesion (p_adj_=0.037), and extracellular matrix organization (p_adj_=0.037). Enriched Hofbauer cell pathways included phagocytosis (p_adj_<0.001), cytokine production (p_adj_<0.001), and macrophage activation (p_adj_<0.001). Enriched pathways in endothelial cells included regulation of cell adhesion (p_adj_<0.001), endothelial cell migration (p_adj_=0.007) and regulation of cell junction assembly (p_adj_=0.002). Enriched stromal pathways included chondrocyte differentiation (p_adj_<0.001), connective tissue development (p_adj_<0.001), and fibroblast activation (p_adj_<0.001). To highlight similarities and differences between cell types, we compared the overlap of enriched biological pathways with an upset plot (**Figure 4)**. Like the number of unique differentially methylated sites, nucleated red blood cells exhibited 270 unique biological pathways. Endothelial cells had the second-greatest number of unique pathways at 57. Of other placental cell types, stromal cells had 29 and Hofbauer cells 8 unique enriched pathways. Immune cells, including Hofbauer cells, shared 29 enriched pathways.

**Figure 4.**
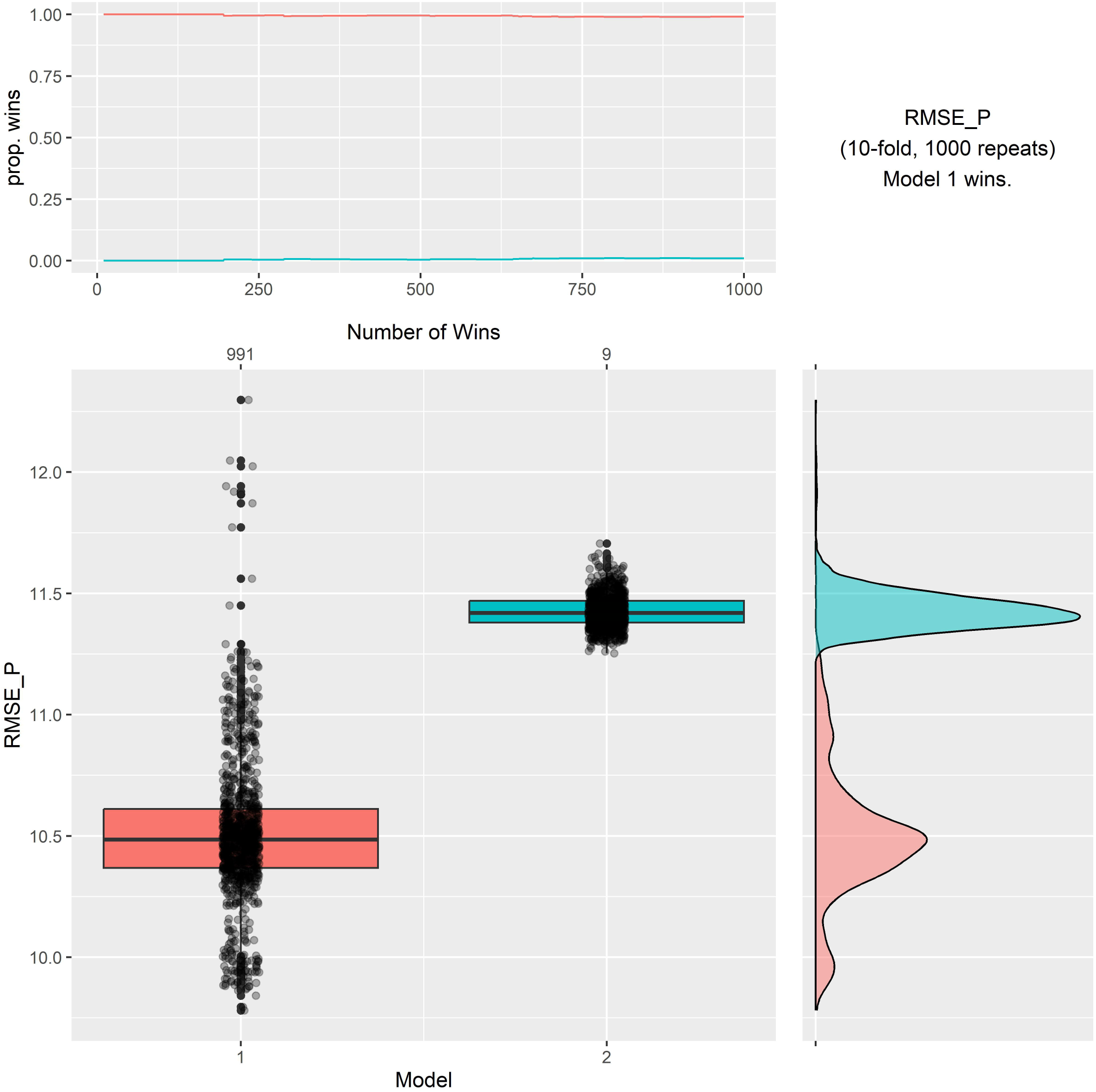
Upset plot of enriched gene ontology biological pathways by placental and immune cell type for the 40 most abundant intersections ordered by intersection size.

### Creation of a robust placental DNA methylation deconvolution reference

To identify cell type-defining DNA methylation sites, we used an existing F-test-based approach [27]. We identified 1,099 unique cell type defining sites (**Figure 5**). Using these sites (**Supplementary Table 3**), we used the robust partial correlation algorithm to estimate the cellular composition of the 35 villous whole tissue samples collected in this study and Yuan et al. [15]. Consistent with placental biology, in whole tissue samples, cell composition median (25^th^ percentile, 75^th^ percentile) estimates were 60.4% (53.3%, 64.2%) syncytiotrophoblast, 17.1% (14.6%, 19.4%) stromal, 8.8% (8.0%, 10.1%) endothelial, 4.5% (3.1%, 5.9%) cytotrophoblast, 3.9% (2.9%, 4.9%) Hofbauer cells, 1.7% (1.3%, 2.0%) nucleated red blood cells, and 1.2% neutrophils (0.9%, 1.6%) (**Figure 6**).Other cell types had median estimates of less than 1%. Monocytes and CD8T cells had median estimates of 0%.

**Figure 5.**
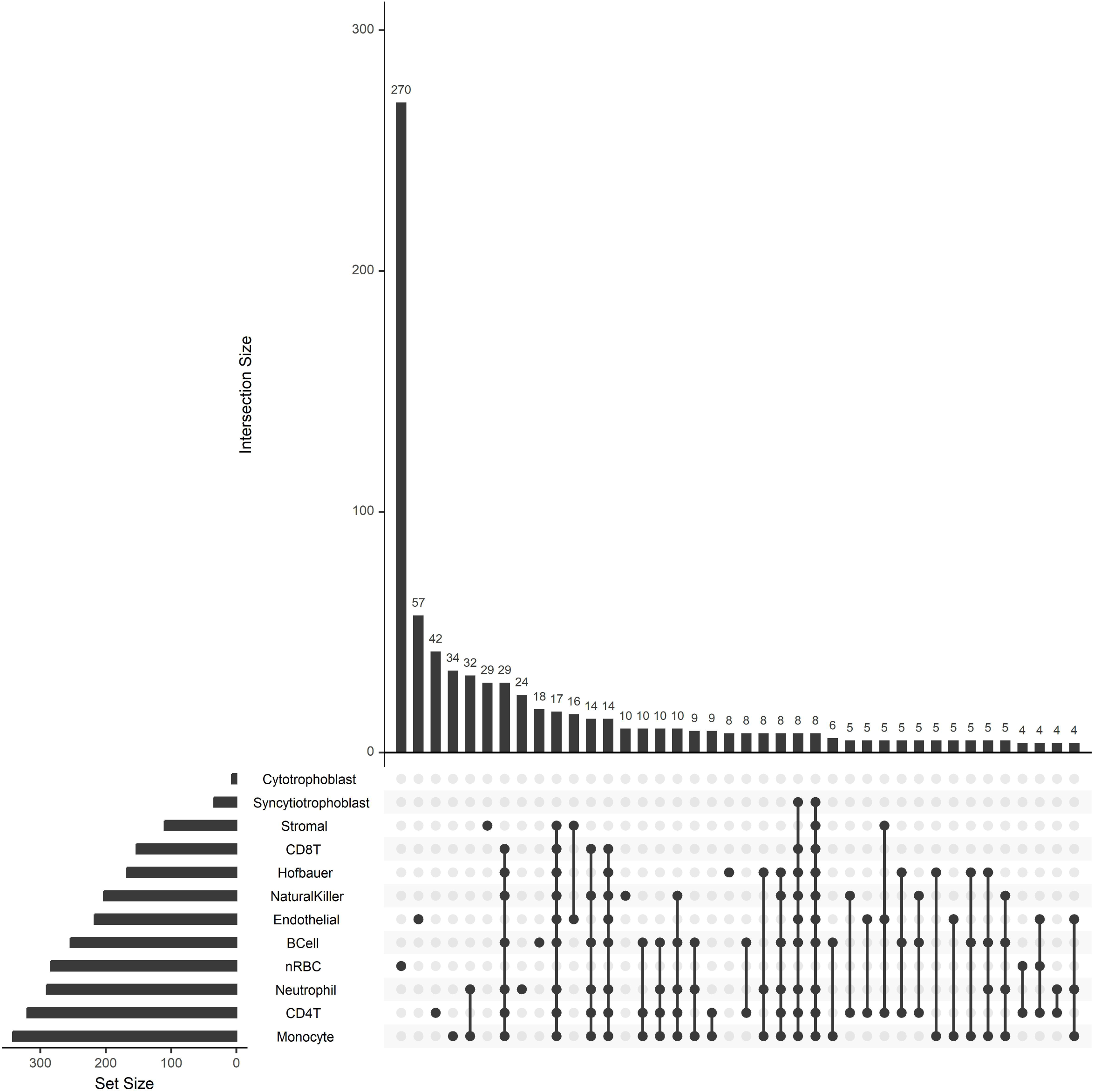
Heatmap of the cell type defining probes used for deconvolution. Color gradient represents percent methylation measure. Abbreviations: nucleated red blood cells (nRBC).

**Figure 6.**
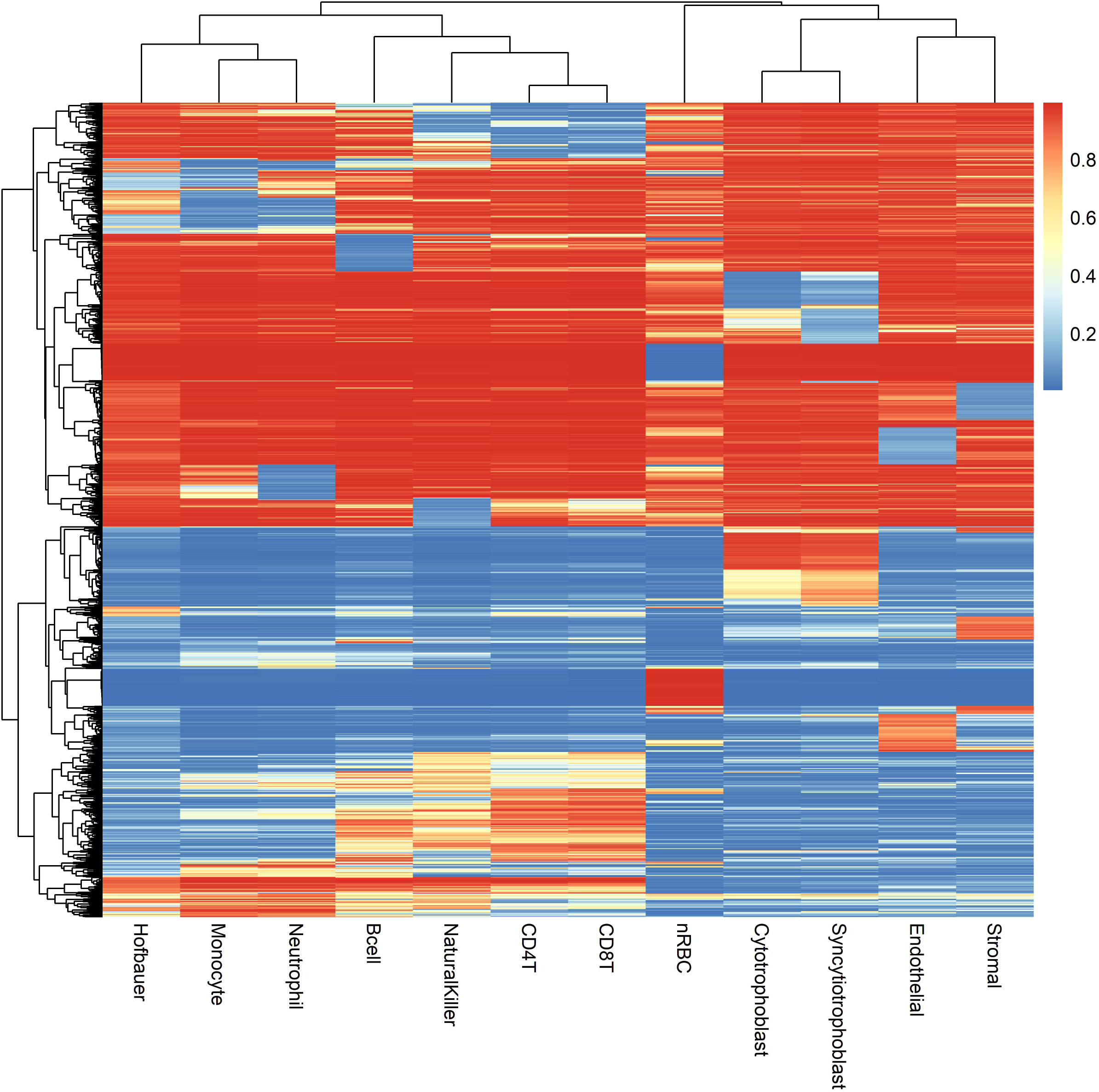
Reference-based cell composition estimates from whole placental villous tissue samples. Abbreviations: nucleated red blood cells (nRBC).

### Comparison between current and previous placental reference panels

We compared the deconvolution performance of our updated placental DNA methylation deconvolution reference against a previously published reference implemented in the planet package [15] with a recently developed deconvolution benchmarking procedure [44]. Importantly, this benchmarking approach does not require direct cell counts, which are currently unavailable. The principal components of whole tissue placental samples provide a prediction target to assess deconvolution performance. We assessed the ability of each deconvolution reference’s ability to predict the first principal component (15.4% variance explained, IQR=21.61) of the site-specific DNA methylation rates of a three-cohort, 204 placental tissue sample dataset in 10-fold cross validation for linear regression models. Our updated deconvolution reference, Model 1, provided a lower root mean square error of prediction (RMSEP) in 991 of 1,000 trials with mean RMSEP 10.51 (SD: 0.30) compared to mean RMSEP of 11.43 (SD: 0.07) for the existing reference, Model 2 (p<0.001) (**Figure 7**). With stricter cell type purity inclusion standards, cytotrophoblast and syncytiotrophoblast samples that clustered more closely with whole tissue samples were excluded. Consequently, our updated deconvolution reference has increased variance in deconvolution performance as tested here. However, this is offset but a much larger decrease in bias that results in a favorable bias-variance tradeoff and overall better performance.

**Figure 7.**
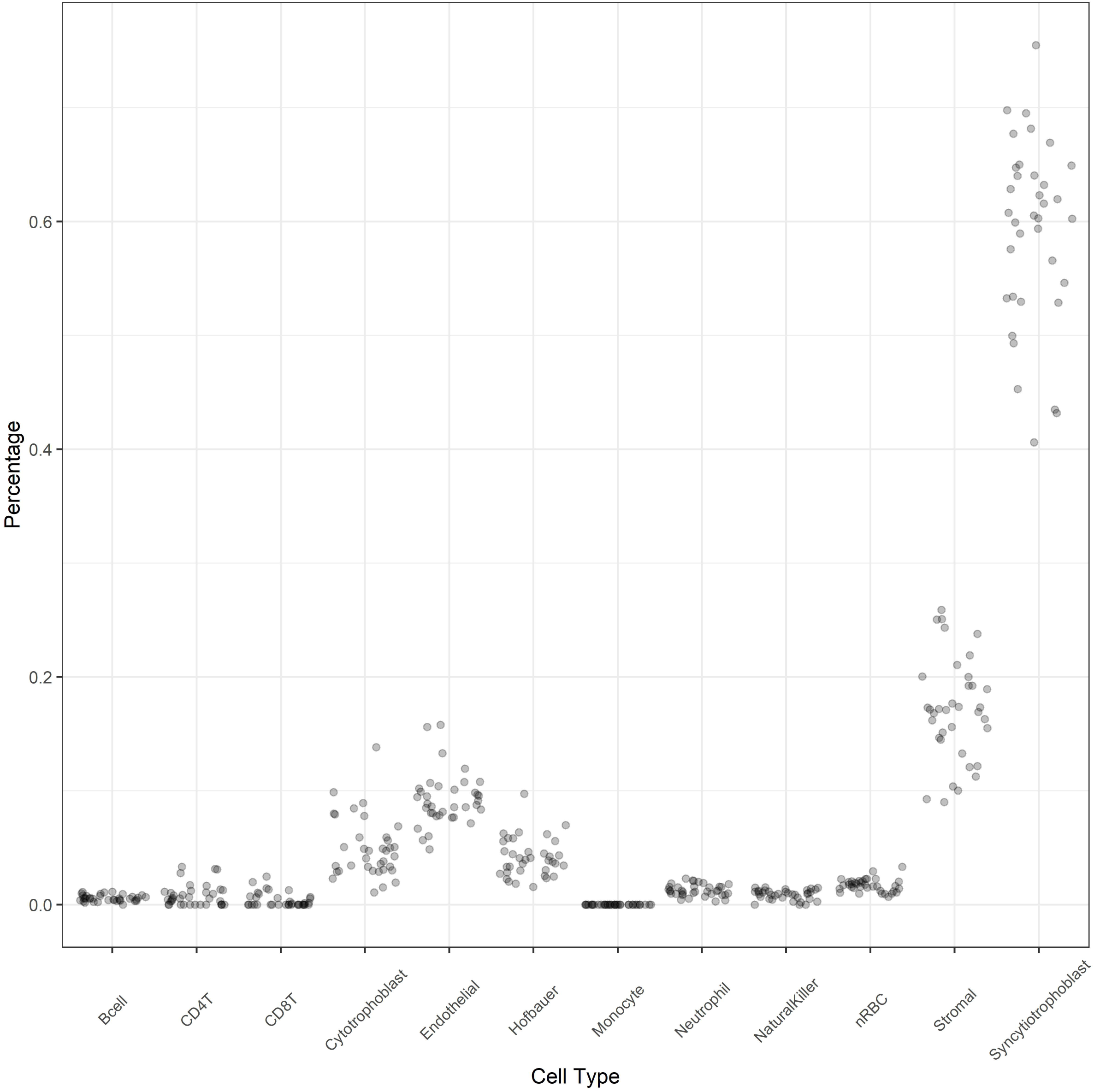
Summary graphs of the root mean square error of prediction in the ability to model principal component one of DNA methylation in 204 placenta dataset GSE232778 for our updated reference (Model 1) versus the existing reference implemented in planet [15] (Model 2) from 1,000 repeats of a 10-fold cross-validation linear regression model with estimated cell composition as the predictor variables.

## Discussion

To provide the most reliable, generalizable, and versatile placental DNA methylation deconvolution reference to date, we generated primary cell type-specific DNA methylation profiles for placental cell types and integrated them with previously published placental cell type and immune cell type samples. We successfully generated primary cell type-specific DNA methylation profiles for placental cytotrophoblasts, fibroblasts, and Hofbauer cells. We uniformly processed and integrated these samples with previously published DNA methylation profiles of placental endothelial cells, Hofbauer cells, trophoblasts, stromal cells, and syncytiotrophoblasts, as well as nucleated red blood cells from umbilical cord blood and B cells, CD4+ T cells, CD8+ T cells, monocytes, natural killer cells, and neutrophils from adult blood. Differential methylation and gene ontology enrichment analyses characterized cell type-defining DNA methylation patterns and their associated biological processes. We confirmed in matched villous tissue samples, that deconvoluted cell type proportions were consistent with expected placental biology. Finally, we showed that our expanded reference, which includes peripheral immune cells, outperforms the only existing placental multi-cell type reference. We propose that this cell type DNA methylation reference panel can robustly estimate cell composition from placental DNA methylation data in epidemiological studies to detect unexpected non-placental cell types, reveal biological mechanisms, and improve casual inference.

The application of deconvolution to address cell heterogeneity in bulk tissue molecular assays and especially its application to the placenta and DNA methylation is a developing field with limited findings. Prior studies were generally limited to one placental cell type and assessed DNA methylation on superseded microarray platforms [12–14]. Yuan et al. represented the most ambitious study to date in developing placental cell type-specific DNA methylation profiles for a deconvolution application [15]. Our independently collected samples were isolated using complementary alternative sorting and antigen marker strategies. Nonetheless, unsupervised clustering resulted in cell type-specific clustering as expected and deconvoluted bulk tissue proportions were consistent with Yuan et al.’s estimates [15]. Despite the alternative cell sorting strategies, cytotrophoblasts clustered with trophoblasts, fibroblasts clustered with stromal cells, and Hofbauer cells clustered with Hofbauer cells.

There are several strengths to this study. We generated placental cell type-specific DNA methylation profiles using established protocols to isolate placental cytotrophoblasts, fibroblasts, and Hofbauer cells [18, 19]. The additional independent samples we collected using magnetic activated cell sorting allowed for sample comparisons with other placental cell types to compare within and across study to identify sample outliers with unsupervised clustering and improve confidence in reference DNA methylation profiles, including using an external resource to show improved lineage identity concordance [32]. By limiting our selection of DNA methylation microarray probes used for deconvolution to only those shared on the 450k and EPIC array platforms, our results will be generalizable to data generated on either platform. The inclusion of nonplacental immune cells in the deconvolution panel allows for the detection of intermingled nonplacental cell types, providing feedback for investigators on placental sampling techniques and identifying potentially cryptic or unexpected cell distribution patterns in study samples. A similar approach has proved beneficial in using DNA methylation data to verify sample-of-origin single nucleotide polymorphism and sex checks in DNA methylation studies to detect sample swaps or contamination [28]. Finally, we leveraged recent developments in reference-based DNA methylation deconvolution benchmarking [44] to evaluate deconvolution performance in a large external dataset of three perinatal cohorts in the absence ground truth cell counts.

There are also limitations to this study. Despite applying an existing syncytiotrophoblast isolation protocol [20], we were unable to characterize syncytiotrophoblast samples that met our strict inclusion criteria. Our ability to confirm deconvolution performance was hampered by a lack of a gold standard reference. This is an ongoing limitation in the field. *In situ* cell counting strategies provide little information about the overall tissue and placental cell sorting techniques are laborious and ill-suited to isolate the syncytiotrophoblast. Further developments in single-cell epigenetics approaches may ameliorate this limitation. Single-nuclei methods may enable more specific characterization of the syncytiotrophoblast [51]. Because current reference-based deconvolution approaches preclude the opportunity to transparently account for batch or confounding effects, particularly when combining data from different studies, our results may be biased by such factors. However, our unsupervised clustering approach highlighted cell type identity as the major driver of DNA methylation variation.

In conclusion, this study developed a robust and flexible DNA methylation reference panel for term placental bulk villous tissue. We provide newly sequenced cell type-specific placental DNA methylation profiles and integrated our samples with existing data, including non-placental cell types that may be present in bulk villous tissue samples. Our independent samples isolated with a complementary approach allowed us to verify and biologically characterize placental cell type-defining DNA methylation profiles. Epigenetics is a promising and increasingly investigated approach to link environmental exposures to biological mechanisms of placental disease and dysfunction. The deconvolution approach developed here advances perinatal epidemiology by allowing investigators to model cell composition in complex study questions to address critical limitations of tissue-level DNA methylation measures.

## Declarations

### Acronyms

APC: allophycocyanin
BSA: bovine serum albumin
DMSO: dimethyl sulfoxide
DNA: deoxyribonucleic acid
EGFR: epidermal growth factor receptor
FBS: fetal bovine serum
GEO: Gene Expression Omnibus
HBSS: Hanks’ balanced salt solution
HEPES: N-2-hydroxyethylpiperazine-N’-2-ethanesulfonic acid
HLA: human leukocyte antigen
nRBC: nucleated red blood cell
PBS: phosphate buffered saline
PE: phycoerythrin
RMSEP: Root mean square error of prediction
TSS200: within 200 base pairs of a transcription start site

### Data availability statement

Data generated in this study are available through the National Institutes of Health Gene Expression Omnibus (accession number pending). This reference panel is available as a Bioconductor R package (link pending). Code to produce the analyses in this manuscript are available through GitHub (https://github.com/bakulskilab).

## Acknowledgements

We thank the participants who provided biospecimens for this study. This research was supported by the National Institute for Environmental Health Sciences (R01 ES025531, R01 ES025574, P30 ES017885, R35 ES031686, R01 ES028802, and P42 ES017198), the National Institutes of Health (NIH), the Office of the Director (UG3 OD023285, UH3 OD023285), NIH and the University of Michigan M+Cubed Program. Dr. Campbell was supported by the National Human Genome Research Institute (T32 HG000040), NIH.

## Author contributions statement

**Kyle Campbell:** Conceptualization, Methodology, Software, Validation, Formal analysis, Investigation, Data Curation, Writing – Original Draft, Visualization **Justin A.**

**Colacino:** Conceptualization, Methodology, Software, Validation, Investigation, Resources, Writing – Review & Editing, Visualization, Supervision, Project administration, Funding acquisition **John Dou:** Software, Validation, Data Curation **Dana C. Dolinoy:** Writing – Review & Editing, Funding acquisition **Sung Kyun Park:** Methodology, Supervision, Writing – Review & Editing **Rita Loch-Caruso:** Conceptualization, Methodology, Resources, Writing – Review & Editing, Supervision, Project administration, Funding acquisition **Vasantha Padmanabhan:** Conceptualization, Supervision, Resources, Writing – Review & Editing, Project administration, Resources, Funding acquisition **Kelly M. Bakulski:** Conceptualization, Methodology, Software, Validation, Formal analysis, Investigation, Resources, Data Curation, Writing – Original Draft, Visualization, Supervision, Project administration, Funding acquisition

## Supplementary Figures and Table

**Supplementary Figure 1.**
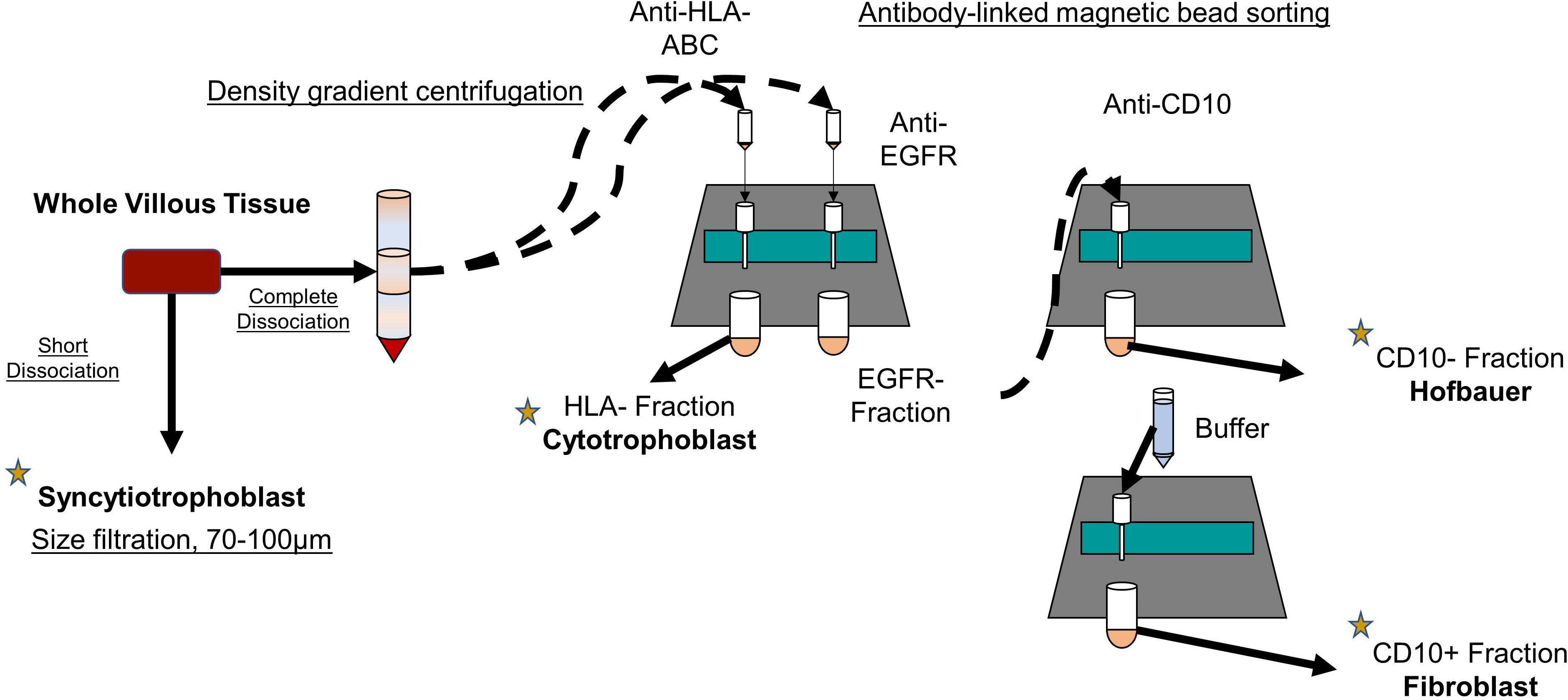
Schematic overview of the magnetic activated cell sorting process.

**Supplementary Figure 2.**
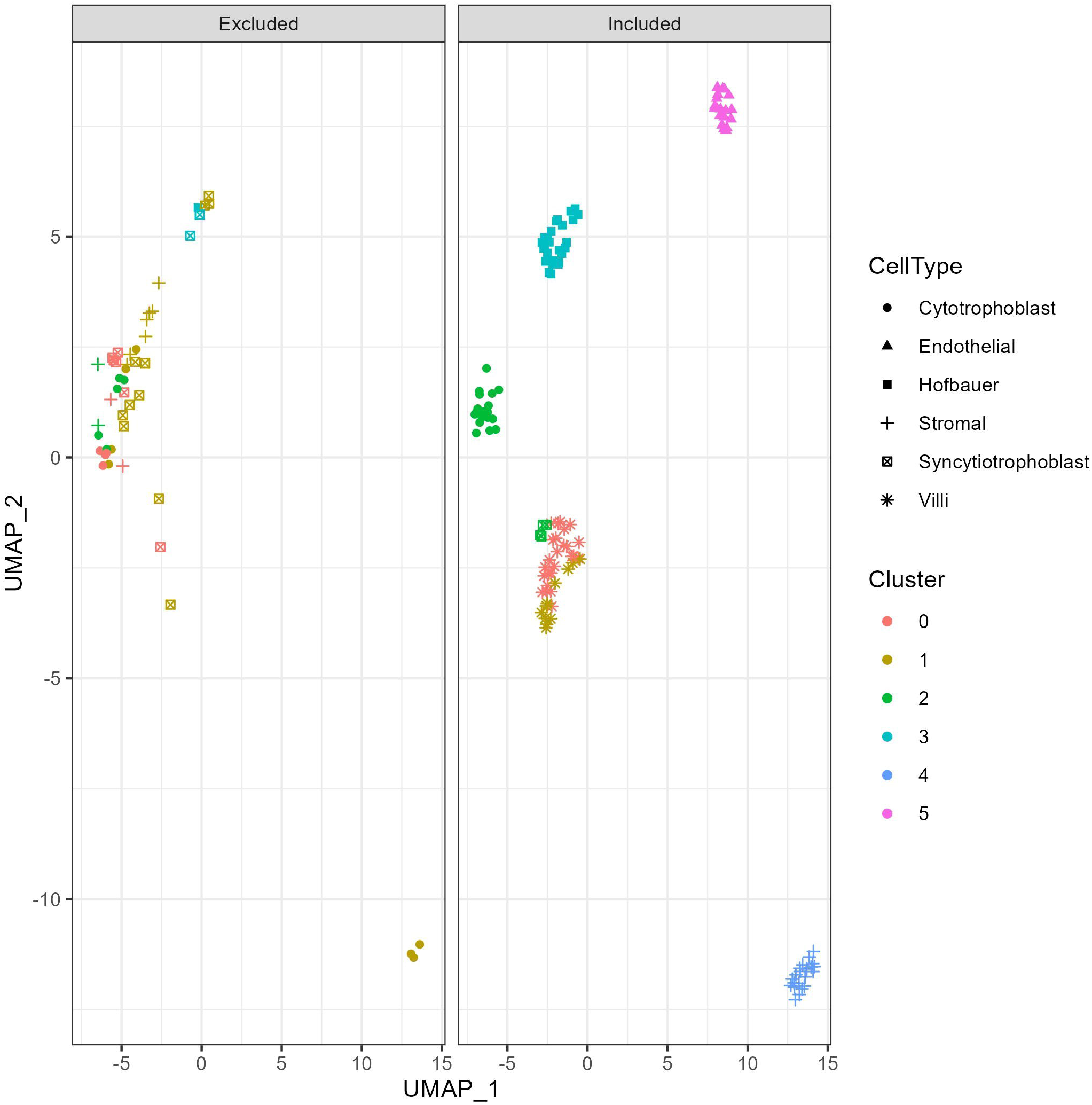
Uniform manifold approximation and projection (UMAP) of DNA methylation data showing how cell type samples that clustered inconsistently with their primary cluster were excluded from downstream analysis.

**Supplementary Figure 3.**
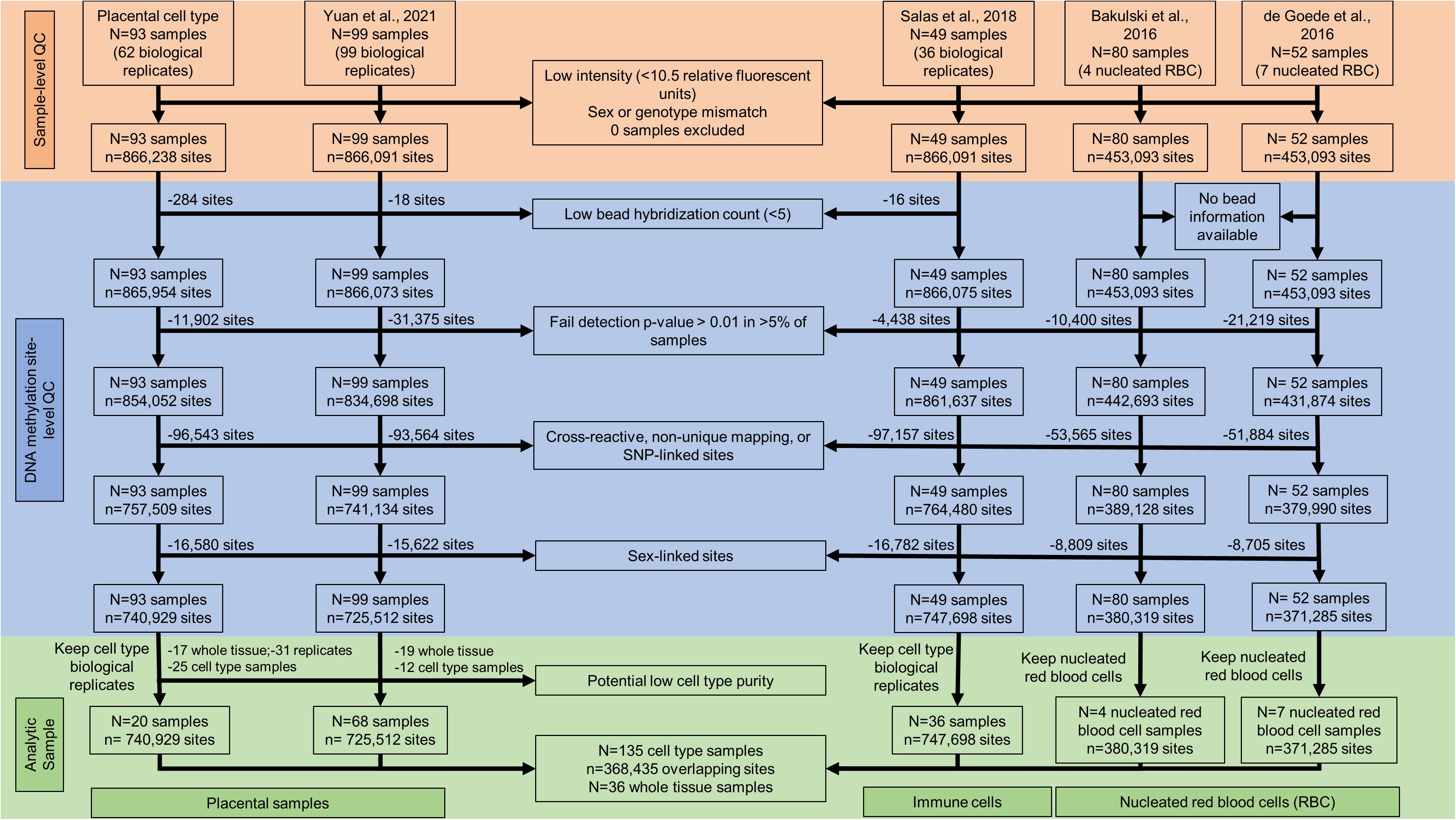
Sample-specific and DNA methylation site-specific quality control inclusion/exclusion flow chart for cell type-specific DNA methylation profiles.

Supplementary Table 1 contains complete differential DNA methylation results comparing the average methylation rate (%), sometimes known as the beta value, in one cell type against the average methylation rate across all 11 other cell types at each of the 368,435 sites assayed. Column headers describe the cell type constrast (cell_type), the DNA methylation site being tested in the Illumina cgXXXXXXXX format (probe_id), the average methylation rate difference (effect_size), the lower 95% confidence interval limit (lower_95_confidence), the upper 95% confidence interval limit (upper_95_confidence), the average methylation rate across all cell type samples (average_methylation_rate), the moderated t-statistic (t_statistic), the p-value (p_value), the false discovery-rate adjusted p-value (fdr_adj_p_value), the B statistics (B_statistic), and a true/false flag for whether the site was considered differentially methylated for meeting our effect size (effect_size > ±10%) and significance (fdr_adj_p_value < 0.001) cutoffs for the analysis (differentially_methylated).

Supplementary Table 2 contains complete differential DNA methylation biological process enrichment results. Column headers describe the cell type contrast (CELL_TYPE), the biological process term ID being tested (TERM_ID), the ontology type (ONTOLOGY), the biological process term being tested (TERM), the size of the term pathway being testing (PATHWAY_SIZE), the number of genes in the pathway that were found to be differentially methylated (DIFFERENTIALLY_METHYLATED_GENES_IN_PATHWAY), the p-value (P_VALUE), the false discovery-rate adjusted p-value (FDR_ADJ_P_VALUE), the names of the genes in the pathway that were found to be differentially methylated (DIFFERENTIALLY_METHYLATED_GENES), and a true/false flag for whether the pathway was considered enriched (ENRICHED).

Supplementary Table 3 contains the deconvolution reference with average methylation rates by cell type that is compatible with the R package EpiDISH’s deconvolution functions as well as other algorithms. The first column describes the DNA methylation site in the Illumina cgXXXXXXXX format (probe_id). Subsequent columns contain the average methylation rate at that site for each of the 12 cell types contained in the reference.

